# qDRIP: Quantitative differential RNA:DNA hybrid immunoprecipitation sequencing

**DOI:** 10.1101/811208

**Authors:** Madzia P Crossley, Michael J Bocek, Stephan Hamperl, Tomek Swigut, Karlene A. Cimprich

## Abstract

R-loops are dynamic, co-transcriptional nucleic acid structures that facilitate physiological processes and cause DNA damage in certain contexts. Perturbations of transcription or R-loop resolution are expected to change their genomic distribution. Next-generation sequencing approaches to map RNA:DNA hybrids, a component of R-loops, have so far not allowed quantitative comparisons between such conditions. Here we describe quantitative differential RNA:DNA immunoprecipitation (qDRIP), a method combining synthetic RNA:DNA-hybrid internal standards with high-resolution, strand-specific sequencing. We show that qDRIP avoids biases inherent to read-count normalization by accurately profiling signal in regions unaffected by transcription inhibition in human cells, and by facilitating accurate differential peak calling between conditions. Finally, we use these quantitative comparisons to make the first estimates of the absolute count of RNA:DNA hybrids per cell and their half-lives genome-wide. Overall, qDRIP allows for accurate normalization in conditions where R-loops are perturbed and for quantitative measurements that provide previously unattainable biological insights.

## Introduction

R-loops are three-stranded nucleic acid structures consisting of a Watson-Crick RNA:DNA hybrid and a displaced single strand of DNA. They typically form during transcription, when nascent RNA hybridizes to its DNA template, and they appear to facilitate certain biological processes while provoking DNA damage in other contexts (1, 2). R-loops are highly dynamic structures that can form in different locations depending on cell type (3, 4) and growth conditions (5). Thus, it is expected that perturbations in growth conditions or depletion of R-loop resolving factors (6, 7) would change their genomic levels and distribution. However, truly quantitative comparisons of R-loop levels between such conditions have so far proven elusive.

A number of recent studies have mapped where R-loops form genome-wide through next-generation sequencing approaches. In the most-widely adopted mapping technique, DNA:RNA immunoprecipitation sequencing (DRIP-seq) (8), the hybrid component of R-loops is directly immunoprecipitated using the S9.6 RNA:DNA hybrid antibody from restriction-digested genomic DNA (9). Various subsequent methods have modified DRIP-seq to increase the resolution by sonication (10–12), to map hybrids to a specific strand (4,10,13), or to capture R-loops in a more native context (14). Increasingly, hybrid mapping has been used to detect regions of change when growth conditions (5) or R-loop processing factors are altered (15–18). In all of these studies, comparisons are made after hybrid signal has been normalized to the total mapped reads from each sample. However, normalizing to total read counts makes a key assumption: that the RNA:DNA hybrid content remains constant between conditions. While this assumption may be appropriate for small perturbations, it is likely inaccurate when the R-loop content significantly changes between samples.

This issue is not specific to RNA:DNA hybrid mapping, and in a range of other next-generation sequencing approaches this type of normalization has been shown to introduce biases, obscure real changes between conditions and even lead to misinterpretations (19–21). Outside of RNA:DNA hybrid sequencing, well-defined internal standards have proven to be a reliable tool to facilitate the accurate comparison of sequencing signal between experimental conditions (20–23). These spiked-in standards or “spike-ins” are introduced during sample preparation, carried through the experimental workflow, and ultimately sequenced to provide an internal normalization factor independent of total mapped reads. In ChIP-seq, *Drosophila* chromatin spike-ins have been used to quantitatively normalize sequencing signal from histone marks on the human genome, revealing a genome-wide depletion of these marks that was not apparent with normalization using total read counts (23). In RNA-seq, synthetic RNA spike-ins reduced bias in quantifying short genes (22) and changed data interpretations compared to standard normalization, thereby reconciling apparently contradictory experimental data (20).

Because RNA:DNA hybrids can be created *in vitro*, pure, synthetic RNA:DNA hybrids are an ideal internal standard to normalize hybrid signal. Here, we describe quantitative differential RNA:DNA immunoprecipitation sequencing (qDRIP-seq), a method combining RNA:DNA hybrid internal standards with a high-resolution version of the strand-specific ssDRIP-seq protocol (13). We show that qDRIP-seq allows for sensitive, high-resolution, strand-specific mapping of RNA:DNA hybrids, and facilitates comparisons in hybrid content between different biological conditions.

## Results

### Design and preparation of RNA:DNA hybrid spike-ins

We sought sequences for the RNA:DNA spike-ins that would serve as effective and orthogonal internal standards for qPCR and next-generation sequencing experiments in human cells. We chose *E.coli* sequences for the spike-ins, which had little homology to the human genome and therefore were unlikely to cause spurious qPCR or sequencing signals. We designed four RNA:DNA hybrid spike-ins (L132, H136, L286, H281), as well as two double-stranded DNA (dsDNA) spike-ins (LDNA, HDNA) and one ssDNA spike-in (ssDNA) as negative internal controls (Supplemental Table S1). In brief, we amplified target regions from purified *E. coli* genomic DNA using primers containing a flanking T7 promoter. RNA was then prepared by *in vitro* transcription using T7 polymerase, purified, and annealed to synthetic complementary ssDNA (Figure 1A). To account for potential biases in sequence recognition with the S9.6 antibody (24), our hybrid and dsDNA spike-ins were designed both with low and high GC-content (Figure 1B). After confirming an RNase-H-reversible size shift in our hybridization product by gel electrophoresis (Figure 1C), we excised the shifted hybrid band and isolated pure hybrids with a gentle gel crushing procedure (25). Ethanol precipitation caused some hybrid dissociation, whereas our purification approach was faster and produced cleaner intact hybrids.

**Figure 1.**
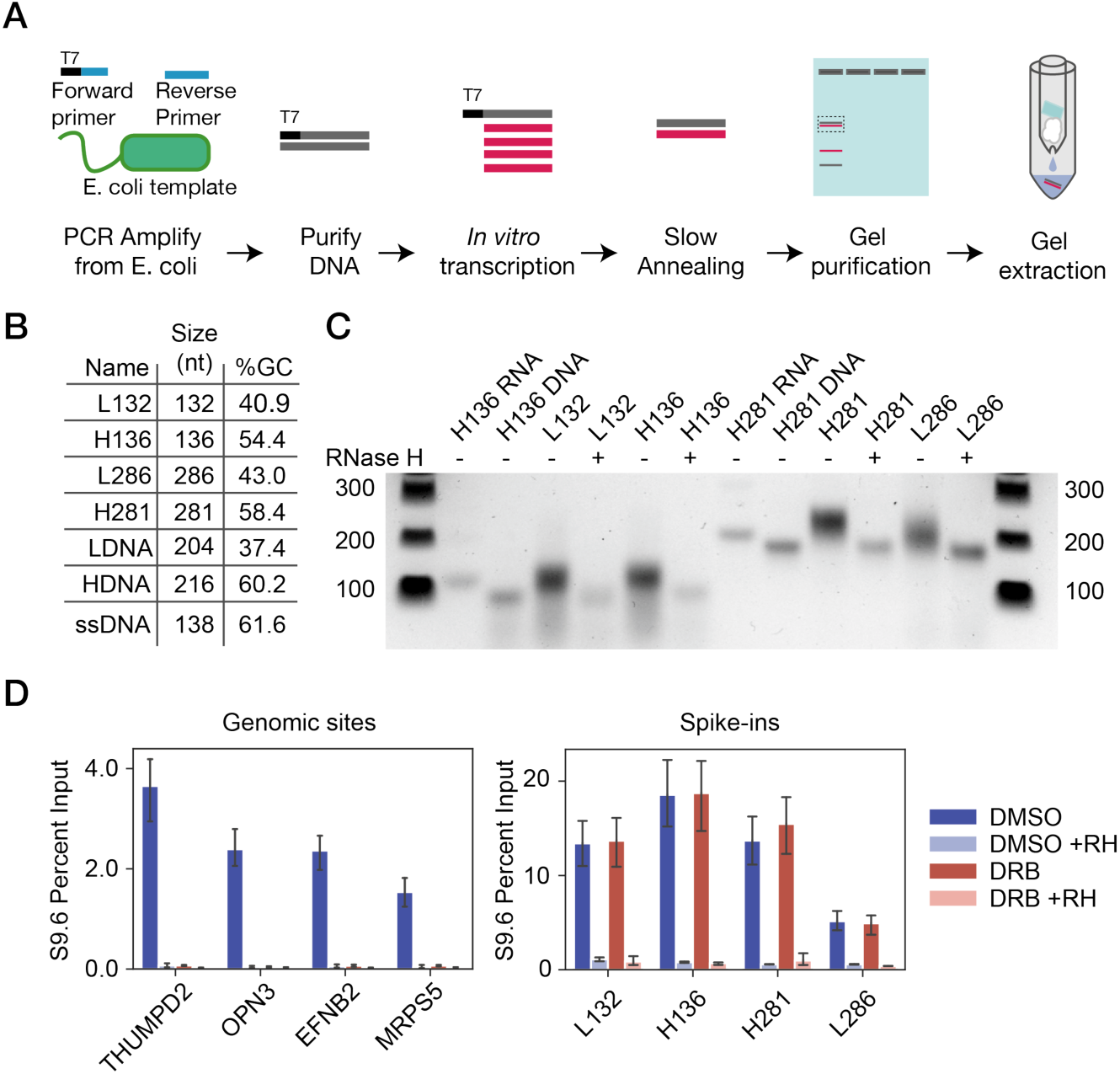
Preparing and evaluating synthetic RNA:DNA hybrids as spike-ins for DRIP. **(A)** Experimental scheme showing how hybrids were synthesized. Briefly, target regions were amplified from *E. coli* genomic DNA with a flanking T7 promoter. RNA was prepared from these templates by *in vitro* transcription, then hybridized to a synthetic ssDNA oligo. Hybrids were purified by agarose gel electrophoresis. **(B)** Length and GC content for the positive and negative control spike-ins used in this study. **(C)** Gel showing RNase H (RH) reversible size-shifts after hybridization of RNA and DNA. **(D)** qPCR of genomic (left) and spike-in (right) hybrids following transcription inhibition with DRB. RNase H treatment demonstrates specificity. Error bars represent 95% confidence interval (CI) of the mean. Results are significantly different as determined by non-overlapping 95% CIs.

We next determined how efficiently our synthetic RNA:DNA hybrids were isolated by immunoprecipitation as compared to genomic hybrids. To best control for technical variation between samples, we introduced spike-ins during the initial cell lysis. Genomic DNA was then extracted from 2 million HeLa cells, fragmented by sonication and DRIP-qPCR was performed. We first confirmed that the spike-ins were efficiently recovered (Figure 1D). Next, we tested whether the spike-ins would remain unchanged after gross hybrid perturbation using 5,6-dichloro-1-β-D-ribofuranosylbenzimidazole (DRB), a potent inhibitor of RNA Pol II transcription that dramatically reduces hybrids at many genomic sites (3, 4). After confirming that DRB reduced transcription outside of the nucleolus by imaging nascent RNA (Figure S1 A, B), we compared changes in levels of genomic hybrids and spike-ins by qPCR. DRB treatment dramatically reduced hybrids at genomic loci, but left the yield of spike-in hybrids unchanged (Figure 1D), indicating that the spike-ins can serve as effective standards for qPCR under strong perturbations of genomic R-loops. In all cases, signal both from genomic and spike-in sites was highly RNase H sensitive (Figure 1D), confirming the purity of the synthetic hybrids and the S9.6 antibody’s specificity for hybrid structures.

### High resolution, strand-specific sequencing of genomic and spike-in hybrids

To sequence both genomic and spike-in hybrids, we adopted an approach similar to ssDRIP-seq which directly sequences the template ssDNA strand of the hybrid (13). As the spike-in hybrids contain only one strand of DNA, we surmised that dsDNA library preparation methods, like those used in DRIP-seq, might not efficiently detect them (8). Sequencing RNA as in RDIP (10) or DRIPc (4) would work in principle, but we sought to avoid RNA-based techniques due to the S9.6 antibody’s off-target affinity for double-stranded RNA (26, 27). We also modified the ssDRIP-seq protocol to fragment the genome by sonication, rather than enzymatic digestion (Figure 2A). This improves resolution and reduces certain biases in hybrid mapping near promoters (12). Additionally, we found that this workflow reduced the necessary number of cells from 10 million to 1-2 million. Only the longer spike-ins (L286, H281) were used for analysis, as sequencing experiments showed that our shorter length hybrid spike-ins (L132, H136) were not efficiently included in sequencing libraries (data not shown). We sequenced IPs from two biological replicates, a matched input and a control IP treated with RNase H prior to pulldown. After adapter trimming, reads were aligned to human genome build hg38, with duplicate alignments removed prior to further analysis.

**Figure 2.**
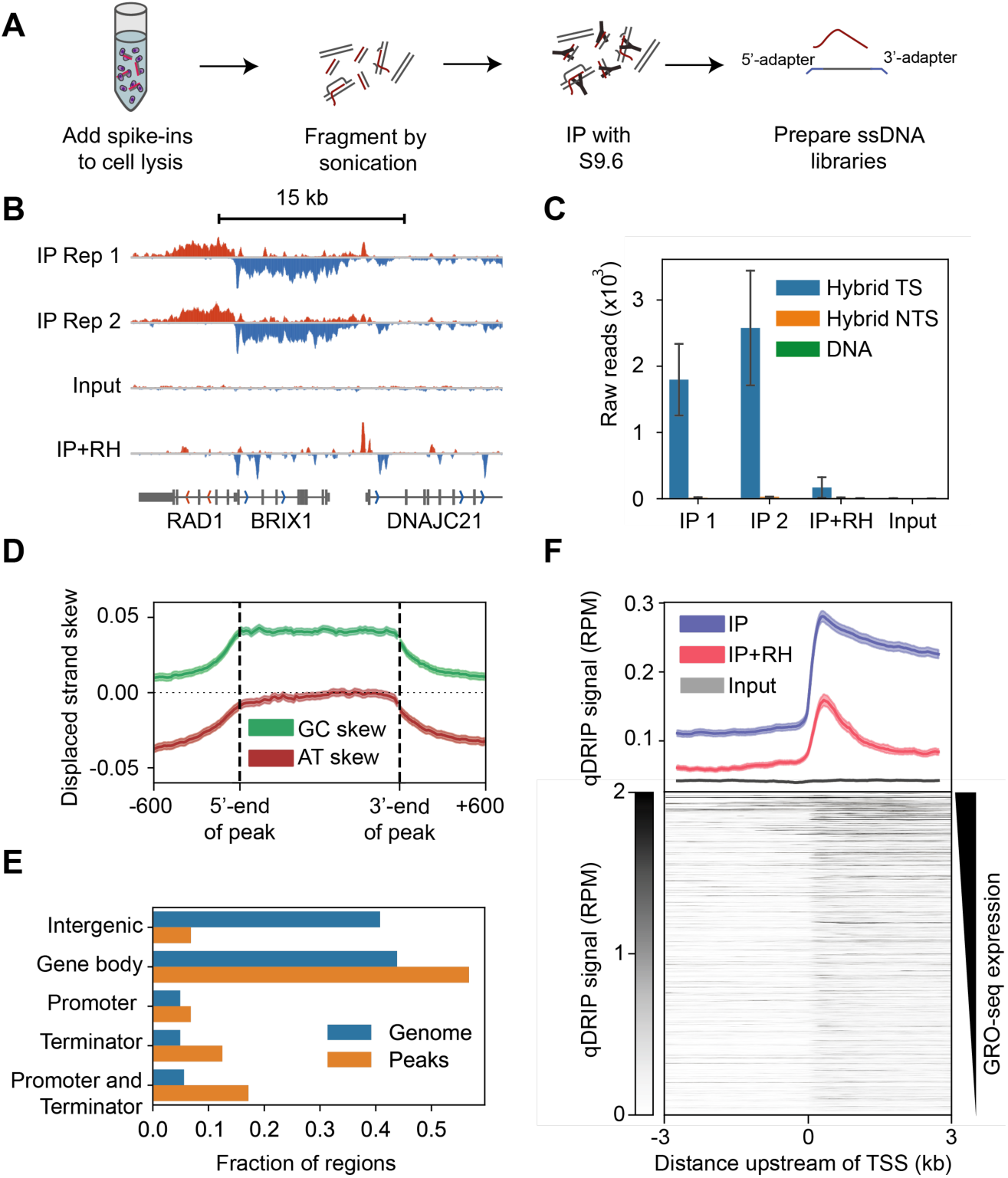
qDRIP provides strand-specific, high resolution RNA:DNA hybrid mapping. **(A)** Schematic of qDRIP experimental process. **(B)** Representative genome browser view of qDRIP-seq signal. From top to bottom: two qDRIP-seq biological replicates, input pooled from replicates, and RNase H digested pool prior to IP. All tracks normalized by reads per million mapped. Negative strand singal in red, positive in blue. **(C)** Read counts from template strand (TS) and non-template strand (NTS) of hybrids, as well as combined from ssDNA and dsDNA negative controls. **(D)** GC (green) and AT (red) skew across coding strand of qDRIP peaks, including 600 bp flanking 5’- and 3’-ends. **(E)** Fractions of qDRIP peaks overlapping noted genomic features. (**F)** qDRIP-seq signal distribution around transcription start sites (TSS).

We found qDRIP-seq signal to derive primarily from the template strand of transcribed areas within the genome, as expected for co-transcriptional RNA:DNA hybrids (Figure 2B). From our spike-ins, we detected significant hybrid signal only from the DNA strand of the synthetic hybrids, but not from the ssDNA or dsDNA controls (Figure 2C). Broad peak calling against a matched input sample identified 65,541 preliminary peaks. To facilitate downstream analysis, Metaplot shows mean of replicates (blue), RNase H sample (red) and input (grey.) Heat map represents qDRIP-seq signal in 10000 highest-expressed genes by GRO-seq (Laitem et al. 2015). we selected 16,895 high-confidence peaks with strong strand bias and RNase H sensitivity, representing 191 Mb (6.3%) of genome space. These values are similar to previous reports of the hybrid-containing fraction of the genome, which have found between five and ten percent of the genome to form hybrids (4, 13). As most R-loops are thought to be 100-2,000 bp in length (6,8,28), and our mean peak size is 11 kb, each peak site likely represents a population average of several individual R-loops, as recently suggested by single-molecule experiments (Malig 2019). Across qDRIP-seq peaks, we found patterns of GC-skew (asymmetry in G content between strands) and AT-skew (asymmetry in A content between strands) as previously reported (Figure 2D) (8, 13). We additionally found qDRIP-seq peaks to be highly underrepresented in intergenic regions and particularly over-represented at the 3’- and 5’-ends of genes (Figure 2E), consistent with other reports (4, 10).

We next asked how qDRIP-seq hybrid signal correlated with known features of the genome. Across promoters, sense hybrid signal was precisely bounded by the TSS (Figure 2F). Beyond the TSS, we confirmed previous reports of hybrids decreasing from the first exon-intron boundary into the first intron (Figure S2A) (14), although we observed a striking increase between the TSS and this boundary. This may reflect a role of the splicing machinery in preventing R-loop formation (29) or increased RNA Pol II pausing at the intron-exon boundary (30). These patterns are consistent with co-transcriptional R-loop formation and demonstrate the high resolution and strand selectivity of our mapping protocol. Beyond the promoter, signal was elevated across the entire gene body with a slight increase at the 3’-end of genes (Figure S2B). Around promoters positive for hybrids, we found that histone modifications associated with active transcription were over-represented compared to an expression-matched set of non-hybrid-containing promoters (Figure S2C), as previously reported for R-loops (4,10,13).

RNase H degrades RNA within hybrids. We therefore predicted that RNase H treatment prior to IP would largely eliminate genomic and spike-in hybrid signal. Indeed, RNase H treatment substantially reduced signal in 88.5% of the area of all called peaks (Figure S3A) and across spike-in constructs (Figure 2C). However, we noticed that some short regions (mean size 519 bp) within qDRIP-seq peaks, particularly near promoters (Figure 2F, Figure S2A,B), were RNase H resistant, and that post-treatment signal was high in some regions lying outside of qDRIP-seq peaks (Figure S3A). These regions were over-represented in GC-skew and G quartets relative to neighboring regions in qDRIP-seq peaks (Figure S3B, C). At promoters, we found RNase H-resistant signal to strongly correlate with the strength of GC-skew and to be mostly limited to regions of strong GC-skew (Figure S3D). This resistant signal may simply reflect incomplete RNase H digestion. However, an intriguing alternative is that these sites form RNA secondary structures which have previously been shown to be resistant to RNase H digestion (31, 32). Because we cannot presently rule out either possibility, for further analyses we excluded RNase H-resistant regions to avoid the possibility of false-positive signal.

### Synthetic spike-ins allow for absolute quantitation of genomic hybrid fractions

Our hybrid spike-ins were prepared *in vitro* and purified, added at levels similar to the genome copy number in HeLa cells, and carried through the entire experimental workflow. Spike-in read counts should therefore be similar to a genomic site that forms R-loops 100% of the time. We thus reasoned that spike-ins could be used to estimate the absolute hybrid frequency at each peak site: that is to say that a site forming R-loops at approximately 5% of all copies in a population of cells would be expected to have approximately 5% of the spike-in read depth. Summing these frequencies throughout the genome, we can therefore estimate the number of hybrids per cell.

To accurately quantify the percent of hybrid-containing molecules at each site, we first estimated the genome copy number at every hybrid peak site by examining input DNA read counts over 10 kb bins. We found a mean DNA content at each locus slightly greater than 3N (Figure S4A), consistent with measurements and known copy number alterations in HeLa cells (Figure S4B) (33). After normalizing the read depth over each qDRIP peak site to its respective genome copy number, we then compared these read depths to the read depth measured over the spike-ins.

We find that the mean qDRIP peak region contains a hybrid at approximately 0.8% of copies (Figure 3A), although some hybrids at housekeeping genes were found to form at rates between 1 and 10%. These percentages are approximately consistent with previous site-by-site measurements obtained by qPCR (34). As comparisons to a pure internal standard also account for dissolution of hybrids during the experimental workflow, these values represent an independent estimate of genomic fractions from qPCR. Using these data along with the assumption that hybrid formation at different sites is statistically independent, we numerically simulated the count of hybrids in each diploid cell. We assigned each peak site as hybrid positive or negative with the probability of its calculated genomic fraction. These probabilistic estimates were used to simulate R-loop content in the genomes of 100,000 diploid cells. Using this simulation procedure, we found the mean unperturbed cell to have approximately 300 R-loops at steady state (Figure 3B). This estimate likely represents a lower bound due to incomplete recovery of hybrids in the sequencing protocol, and the strong possibility that more than one distinct R-loop structure may form within a single genomic peak site.

**Figure 3.**
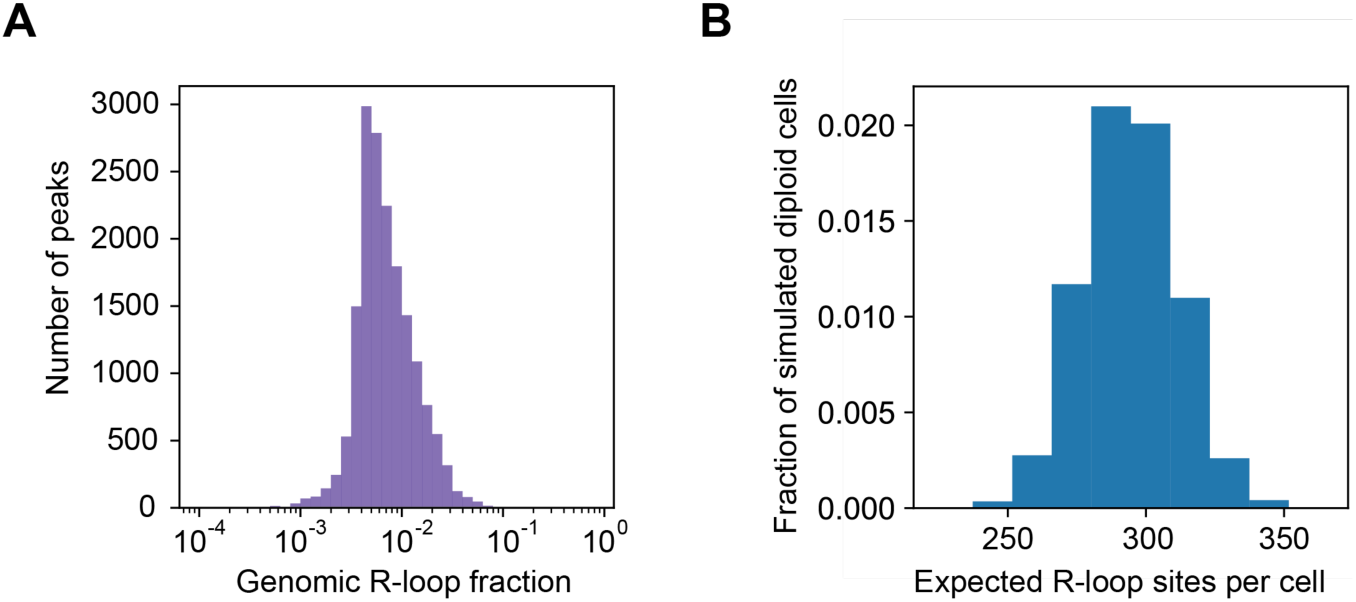
Absolute quantitation of genomic hybrids using the spike-in. **(A)** Histogram of estimated maximum genomic hybrid frequencies across consensus qDRIP peaks. Data is pooled from mean of two biological replicates. **(B)** Histogram of distribution of R-loop frequencies in diploid cells obtained from numerical simulation on genome-frequency data.

### Synthetic RNA:DNA spike-ins allow for accurate hybrid signal normalization across R-loop perturbations

Having established a method to sequence genomic and synthetic RNA:DNA hybrids, we next evaluated the spike-ins under experimental conditions where R-loop formation is severely perturbed. As a proof of principle, we chose to examine hybrid levels in the presence and absence of the RNA Pol II transcription elongation inhibitor DRB, which should acutely perturb R-loop formation at predictable sites. In particular, we expected that DRB treatment would reduce hybrids in places where DRB inhibits transcription, such as downstream of the pause site in Pol-II-transcribed genes, but not within non-pol-II genes such as ribosomal DNA. As DRB would reduce the total number of hybrid-containing sites on the genome, we also expected that raw read counts across remaining areas would be correspondingly increased. If normalizing by total read counts, this would lead to an artifactually high signal at these regions after DRB treatment relative to controls. A similar pattern of overestimation has previously been observed for nascent transcription by GRO-seq after DRB treatment (35).

We thus performed qDRIP-seq in HeLa cells treated with 100 µM DRB for 40 minutes, a time-point previously shown to reduce but not eliminate hybrids at a subset of human genes (4). As expected for inhibition of transcription elongation, treatment with DRB severely affected hybrid formation (Figure 4A), reducing hybrid signal within the 5’-ends of long genes while retaining hybrids lying far downstream from promoters. As short DRB treatment is not expected to affect transcription from non-RNA Pol-II-transcribed genes, these genomic regions effectively act as a natural internal standard that can be used to evaluate how effectively synthetic spike-ins allow accurate normalization between samples (35). At the polymerase-I-transcribed 18S and 28S rDNA subunits, we confirmed by qPCR that hybrid signal remained unchanged (Figure S5A). Using read counts over annotated Pol I and Pol III genes, we found that synthetic spike-in hybrids significantly reduced the extent to which signal at these regions was overestimated (Figure S5B). Similarly, in metaplots of qDRIP-seq read density of Pol I genes, we found that normalizing with read counts overestimated the hybrid signal under DRB treatment. By contrast, normalizing using spike-ins roughly equalized signal between the control and DRB treatment conditions (Figure 4B, Figure S5C).

**Figure 4.**
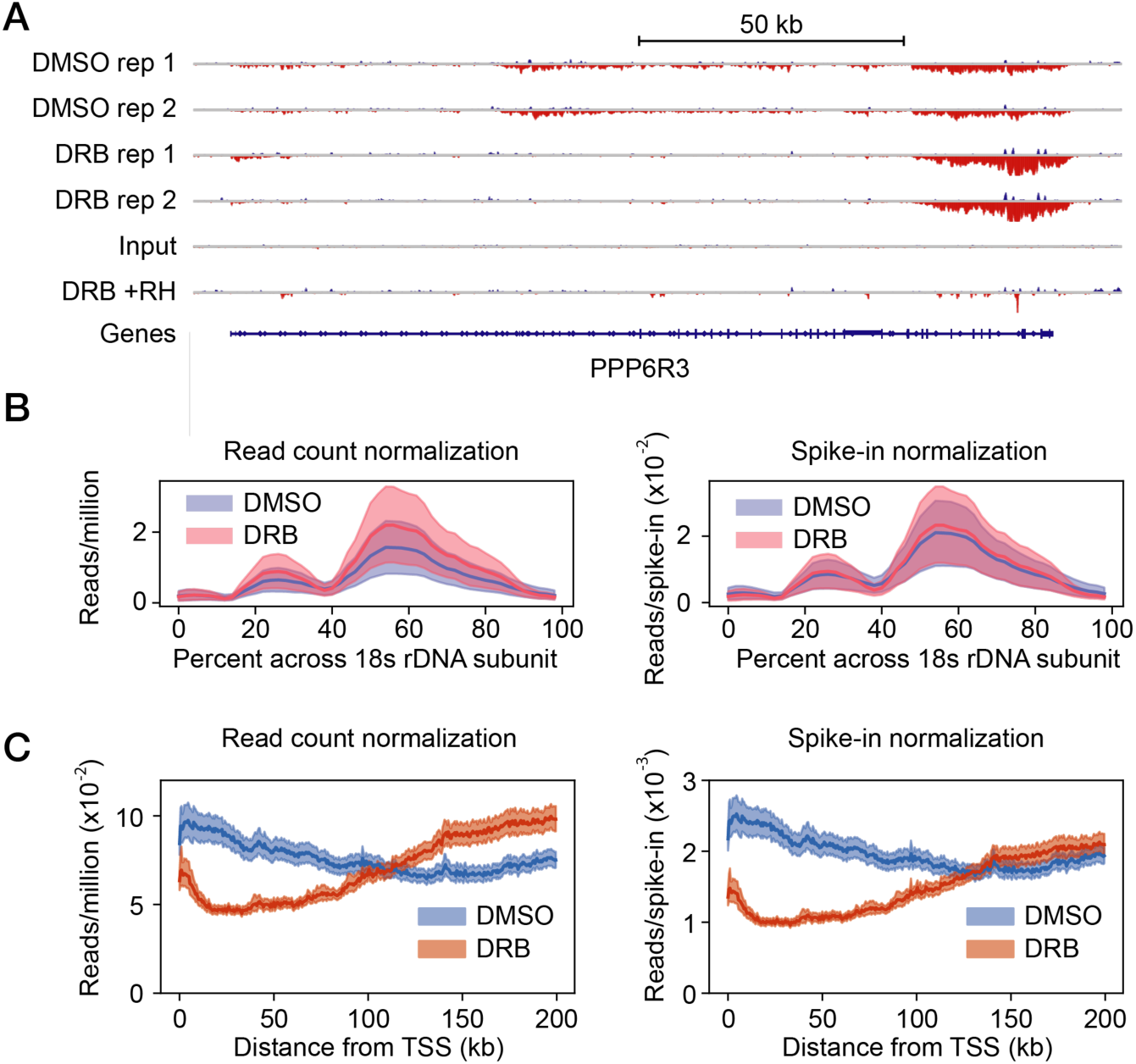
qDRIP allows for effective normalization when R-loops are acutely perturbed by transcription inhibition. **(A)** Genome browser view taken from long gene showing effects of DRB on hybrid formation. In order from top to bottom, tracks shown are 2 biological replicate control (DMSO) IP samples, 2 biological replicate DRB-treated IP samples, overlaid pooled inputs from the control and DRB samples, a pool of the DRB replicates pre-treated with RNase H prior to IP, and a track showing genes. All tracks are normalized by reads per million mapped. Red indicates negative strand signal, while blue indicates positive strand signal. **(B)** Metaplots of DMSO (blue) and DRB (red) IP signal over length of Pol I-transcribed 18s rDNA unit normalized using total read counts (left) or mean spike-in counts (right). As before, bands represent 95% CI of mean read signal. **(C)** Metaplots of DMSO (blue) and DRB (red) IP signal over the first 200 kb of all expressed genes over 200 kb in human genome, normalized using total read counts (left) or spike-in read counts (right)

As a CDK9/PTEF-b inhibitor, DRB blocks transcription by preventing engaged RNA polymerases from transitioning to processive elongation after the 5’-pause site (36). However, DRB does not affect polymerases that have cleared this transcriptional checkpoint. We thus expected that transcription at the 3’-end of long genes should be largely unaffected by DRB treatment. Using publicly available GRO-seq data collected from DRB-treated and control samples (35), we confirmed that nascent transcription remained mostly unchanged at these sites after 30 minutes of treatment using the same dose of DRB (Figure S5D). We reasoned that R-loop formation should also remain largely unchanged in these regions after DRB treatment. As hypothesized, normalization of hybrids with mapped read counts showed a marked increase at the 3’-end of genes (Figure 4A, C). In contrast, the signal was equalized at these regions using spike-in read counts, consistent with nascent transcript levels and the known biology of DRB (Figure 4C). Interestingly, the DRB hybrid signal was equivalent to that in the control approximately 130 kb downstream of the TSS. This is consistent with measurements of transcriptional elongation rates in HeLa cells which suggest that an average polymerase would have travelled 135 kb in 40 minutes (37). Thus, we find that normalization by total read counts consistently overestimates hybrid levels in regions that should be unaffected by DRB, whereas normalizing with spike-ins brings these regions close to parity.

### Hybrid spike-ins facilitate accurate differential peak calling

Having shown that synthetic spike-in hybrids could effectively normalize qDRIP-seq signal across genome features that are unaffected by DRB treatment, we next asked whether spike-ins could facilitate unbiased discovery of differential hybrid-containing regions. In most sequencing experiments, regions where IP signal changes between conditions are not known *a priori*, and are generally discovered through differential peak calling. We thus performed differential peak calling between our DMSO and DRB samples over the union of peaks from both conditions. We selected differential regions between the samples at a 1% false discovery rate, and further filtered these calls to include only strand-specific and RNase H-sensitive qDRIP-seq peaks.

With spike-ins, we found 94 hybrid sites that were significantly induced and 4,812 sites that were significantly repressed after DRB treatment (Figure 5A). Using qPCR, we confirmed that regions called as significantly down (Figure S6A) and up (Figure S6B) in DRB were indeed significantly altered. Without spike-in normalization, we found that 754 (4.5%) peaks were called as increased in hybrid content under transcription inhibition, an increase of 660 peaks (Figure 5A). We additionally found 394 fewer peaks called as significantly down with read-count normalization compared to spike-in normalization. Given the co-transcriptional nature of R-loops, we expect that most regions should decrease in hybrid formation after DRB addition. Therefore, we suspected that many regions called as increased only in the read-count normalization were likely false positives, and regions that failed to be called as down in the read-count normalization were likely false negatives.

**Figure 5.**
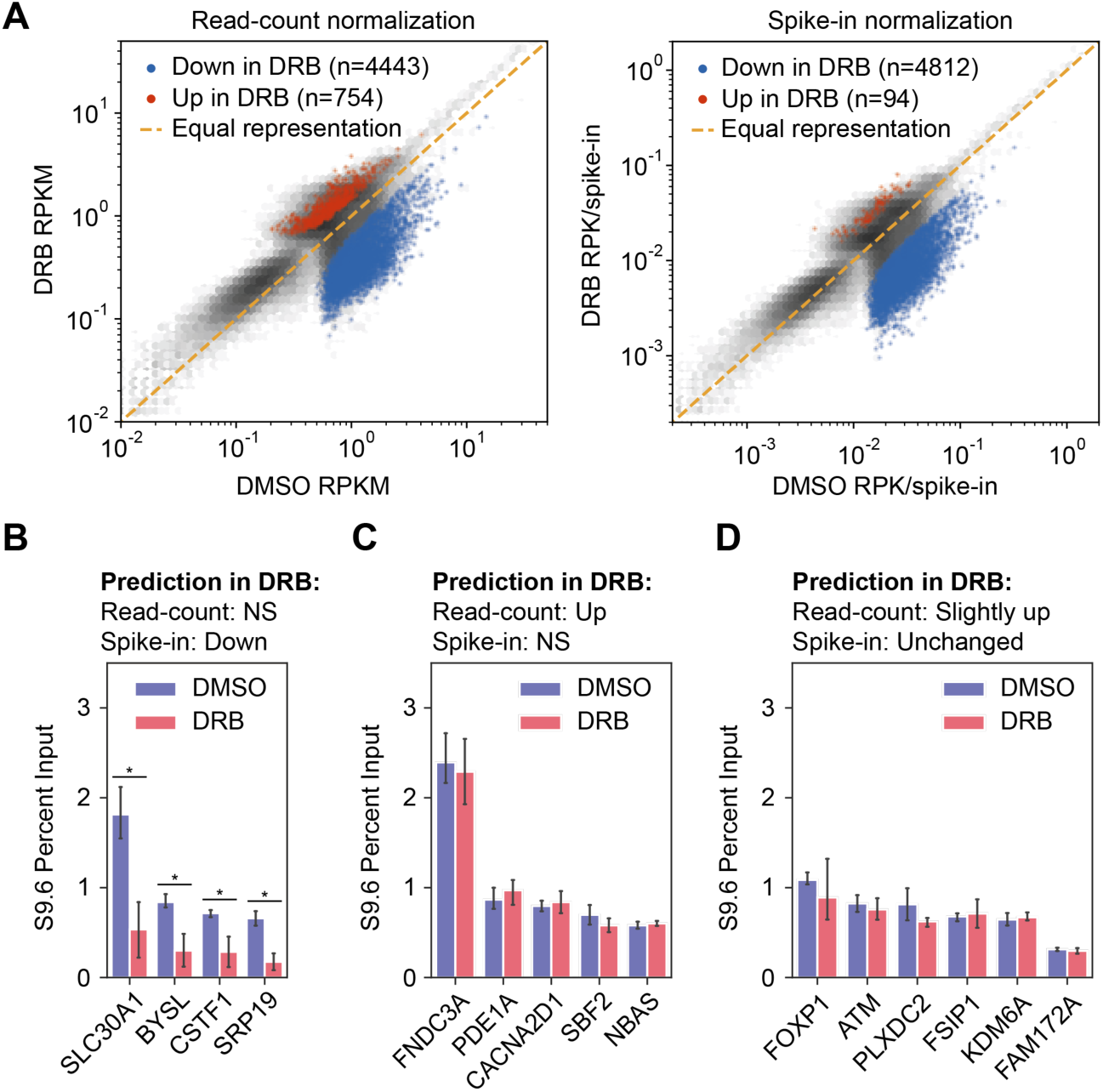
Spike-ins facilitate accurate differential peak calling. **(A)** Comparison of DESeq2 calls normalized using total read counts (left) or DESeq2 calls normalized using spike-in read counts (right). Using a cutoff of 0.01 for the FDR corrected p-value, sites significantly increased in the DRB are highlighted in red, and sites significantly decreased in the DRB highlighted in blue for both normalization methods. **(B)** qPCR measurements over genes called as significantly down by spike-in normalization, but non-significant by total read counts. For all qPCR measurements, error bars represent 95% CI of mean value. Results are significantly different (marked as *) as determined by non-overlapping 95% CIs. **(C)** qPCR measurements over regions called as non-significant by spike-in normalization, but significantly up by read-count normalization. **(D)** qPCR validation of regions called as having fold-changes close to zero by spike-in normalized calls, but slight increases in the DRB by read-count calls.

To test whether these classes of peaks actually represented false negative and false positive calls, we examined changes at a few representative peaks from each class by qPCR. At the peaks that were significantly decreased after DRB treatment only with spike-in normalization, qPCR measurements confirmed a significant decrease (Figure 5B). Thus, normalizing our differential peak calls with spike-ins strikingly increased the sensitivity of these calls. We additionally tested the specificity of our calls by examining regions called as increased by read-count normalization, but unchanged by spike-in normalization. By qPCR, these regions had no significant differences (Figure 5C), again confirming that spike-ins properly normalized differential calls. Finally, we selected regions that were predicted to have almost no change between our conditions as called by spike-in normalization, but that would be predicted as slightly (1.2-fold) increased in the DRB sample by read-count normalization. Measuring these regions by qPCR, we found that levels were roughly equal between conditions and did not show any consistent bias towards greater signal in DRB (Figure 5D). This indicates that spike-in normalization is quantitatively predictive of changes in regions that are not significantly altered between conditions. Taken together, these results show that normalization with spike-ins substantially increases the sensitivity and specificity of our differential peak calls, and confirms the importance of using internal standards to discover regions that change in hybrid content between biological conditions.

### Determination of hybrid lifetimes

Having shown that our synthetic spike-in hybrids could accurately normalize between control and DRB-treated samples, we leveraged the spike-ins as a tool to carry out a natural experiment to estimate R-loop lifetimes genome-wide. Because DRB inhibits new transcription but does not halt actively elongating RNA Pol II, it does not instantaneously halt transcription across the entire gene body. Instead, it leaves a final “front” of transcription from the last polymerases that began elongation before DRB treatment. Using gene-specific rates of transcription previously determined by 4sUDRB-Seq (37) and the length of our DRB treatment (40 minutes), we can estimate the time that DRB has prevented new transcription at a given region in the gene body. Thus, every gene effectively contains a natural time-course of transcription inhibition (Figure 6A).

**Figure 6.**
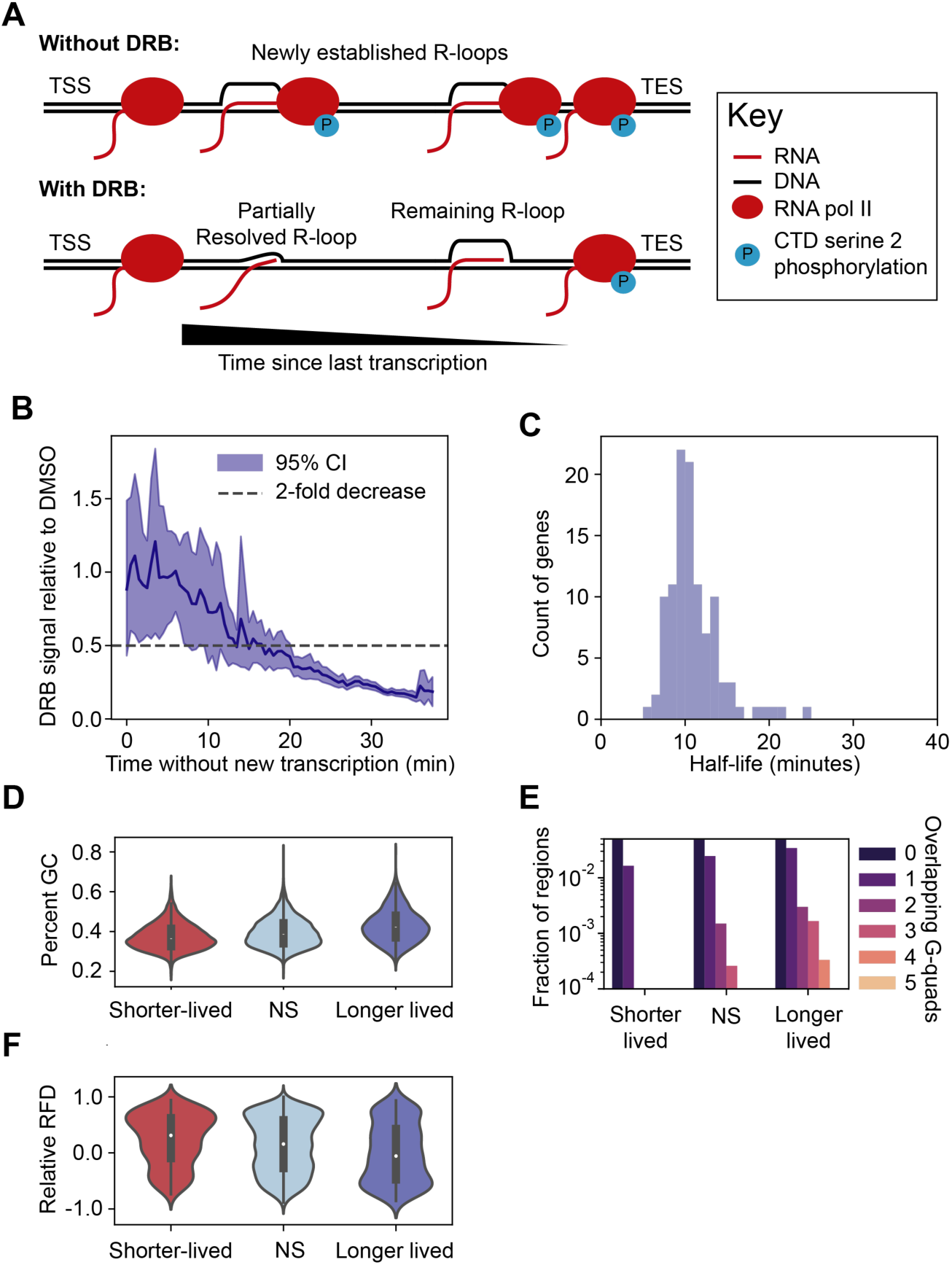
R-loop lifetimes. **(A)** Schematic of transcription with and without DRB. **(B)** Ratio of DRB to control signal in RNase H-sensitive peaks, over estimated time without transcription. Error bands are 95% CI of the mean. Horizontal dotted line indicates a 2-fold decrease in DRB signal. **(C)** Distribution of half-lives assuming first-order decay. **(D)** GC content across 500 bp regions with shorter, longer or close to average lifetimes (p = 2.5e-143, Kruskal-Wallis test). **(E)** G-quadruplex counts over same regions as (C) (p = 2.7e-7, ANOVA on Negative Binomial regression, likelihood ratio test) **(F)** Relative replication fork directionality (RFD) to transcription over same regions as (C), where 1 represents fully co-directional and -1 represents fully head-on (p = 3.5e-12, Kruskal-Wallis test).

Without new transcription, R-loop levels should decrease over time from endogenous cellular R-loop resolution activities. Indeed, previous qPCR experiments found that promoter hybrid levels diminish with a half-life of approximately 10 minutes during a DRB treatment time-course (4). To determine R-loop lifetimes genome wide, we first restricted our search to high confidence qDRIP-seq peaks. Within these regions, we determined the extent to which DRB reduced hybrid signal, and compared this reduction to the inferred time without new transcription. At short inferred time-points of inhibition, we observed parity between the hybrid content in DMSO and DRB (Figure 6B), consistent with our previous findings at the ends of long genes (Figure 4D). However, as the gap between the effective site-specific inhibition of transcription and our hybrid measurements widened, we found a striking reduction in signal. Hybrid levels declined to approximately 20% of what is obtained in control samples, with a half-life that broadly fell between 10 and 20 minutes (Figure 6B).

We next sought to more precisely characterize the individual lifetimes of specific hybrids. Previous measurements from DRB time-courses using qPCR (4) have found hybrids to be resolved in a manner roughly consistent with first order (exponential) decay. To generalize this pattern, we compared linear and exponential fits to decay times, to determine whether an exponential model could better explain our observations than a simple alternative. As we only had one time-point of DRB inhibition, we fit each curve to time-points collected from a single gene, reasoning that sites within a single gene would likely be resolved by similar co-transcriptional mechanisms. In 224 (85%) of these genes, we found that exponential models fit the data better as measured by Pearson’s R^2^ (Figure S7A). Furthermore, the generally high quality of these fits confirmed that site-to-site variation in lifetimes was fairly low across these gene bodies. Among the 107 genes fitted extremely well (R^2^ > 0.95) by an exponential model, we found that 88% had hybrids with mean half-lives between 7 and 15 minutes, with a mean of 11.0 minutes (Figure 6C). These values suggest that previous measurements at selected promoters (4) generalize across the genome.

Having characterized the variation in hybrid lifetimes between genes, we next sought to identify factors that might influence lifetimes within genes. We therefore examined individual 500 bp windows of hybrid-forming sequence that were resolved either significantly slower, faster, or roughly equivalently to the average. Strikingly, hybrid lifetimes strongly correlated with GC content, with shorter-lived hybrids having lower average GC-content and more stable hybrids having higher GC content (Figure 6D). We additionally found that an increase in G-quadruplex forming sequences correlated with longer hybrid lifetimes (Figure 6E). Interestingly, we observed that longer-lived hybrids appeared to be associated with more sites predicted to participate in head-on replication-transcription collisions (Figure 6F), consistent with measurements of higher hybrid content in these regions (38). We did not observe any consistent trends for GC-skew or AT-skew (Figure S7B, C), although we did notice that longer-lived hybrids tended to have more extreme values of both. Lifetimes also did not strongly correlate with sense transcription, suggesting that total transcription may influence hybrid levels (Figure 2F) through formation but not resolution (Figure S7D).

With genome-wide estimates of both the hybrid count per cell and the mean lifetimes of these hybrids, we next estimated the rate at which cells resolve hybrids. A half-life of 11 minutes with first order decay implies that 6.3% of cellular hybrids are turned over every minute. Using our previous result that cells contain approximately 300 hybrids at steady state, we calculated that 19 hybrids are resolved every minute, for a total of 27,000 per day. As a frame of reference, this is roughly double the rate of depurination in human cells (39). Altogether, these results underscore the power of absolute quantitative measurements as provided by qDRIP.

## Discussion

A persistent challenge for the R-loop field has been accurate comparison of hybrid levels between conditions where R-loops are perturbed. More generally, normalization using the conventional approach of total read count assumes that total signal remains unchanged between samples in next-generation sequencing experiments. Sequencing experiments under conditions that strongly decrease or increase signal at a subset of genomic sites will therefore inherently suffer from over- or under-estimation of signal at unchanged sites, respectively. Spiked-in standards have been shown to correct these biases, revealing changes between conditions that were otherwise obscured, and preventing misinterpretations (19–23), but this approach has not hitherto been developed for R-loop mapping.

Here we describe qDRIP-seq, a method that combines stranded, high-resolution hybrid sequencing with synthetic RNA:DNA hybrid spike-ins for cross-condition normalization. We first show that our sequencing procedure recognizes hybrid-containing sites generally consistent with known biology (Figure 2). We additionally use the spike-ins to make absolute estimates of the genomic R-loop fraction at different genomic sites, and used these estimates to model the count of hybrids in an average cell (Figure 3). In cells treated with the Pol II transcription elongation inhibitor DRB, we also show that normalization using total read count overestimates hybrid signal at non-pol II transcribed genes and the ends of long genes. By contrast, normalization using synthetic RNA:DNA hybrid standards corrects these biases (Figure 4). Finally, we find that the use of hybrid spike-ins reduces the false-positive and false-negative rates of differential hybrid peak calling between control and DRB-inhibited conditions (Figure 5).

Here, we provide the first data of hybrid lifetimes at a genomic scale (Figure 6). Measuring kinetic off-rates as opposed to thermodynamic steady state levels could prove to be a powerful new approach to study R-loop biology. For example, hybrid lifetimes would be expected to increase after depletion of R-loop processing factors. Thus, identifying the specific sites where hybrid lifetimes increase could reveal where these factors act. As our lifetime estimates critically rely on the ratio of DRB treated to control R-loop signal, spike-in normalization was crucial for accurate results because normalizing by total read count overestimates hybrid signal at the ends of genes. Interrogating lifetimes genome-wide allowed us to discover that high-GC content and G-quadruplex formation, but not high transcription or nucleotide skew, correlate with longer R-loop lifetimes on the genome. Where sufficient data was available across genes to make lifetime estimates, we found hybrid levels at most genes diminish exponentially with a half-life of approximately 11 minutes. While many factors may influence R-loop lifetimes, the observed exponential decay implies that R-loop resolution does not increase when R-loops are depleted across the genome. This implies that R-loop resolution is not rate-limiting at steady state.

Our estimate that mammalian cells must resolve on the order of 27,000 R-loops per day brings into clear focus the substantial resources that cells must invest to turn over R-loops. Even if 99 percent of R-loops occur in contexts where they are benign, this would leave 260 potential detrimental events per day, a rate comparable to serious events such as DNA base damage (39). If R-loops only cause damage in specific contexts as some recent studies suggest (40, 41), it would be interesting to understand what necessitates this speedy resolution even outside of these contexts. Additionally, this high rate of resolution may partially explain the large and diverse set of pathways required for R-loop processing and resolution, which appear to occur quickly in many different genomic contexts (6, 7).

While this study provides valuable quantitative insights, there is still room for improvement in the method as described. In particular, our conclusions are based on counts from only two spike-in sequences. Because we prepared these spike-ins using a long synthetic DNA oligonucleotide, the cost to produce a large library such as the one provided by the External RNA Controls Consortium (ERCC) (22) could be substantial. Two spike-ins were sufficient to correctly normalize between DMSO and DRB, but additional spike-ins might provide more statistical certainty that could help to normalize conditions with more subtle R-loop perturbations. Nevertheless, we have demonstrated in principle the value of using spike-ins for R-loop mapping genome-wide.

In summary, qDRIP-seq provides high-resolution, strand-specific maps of RNA:DNA hybrids, and allows for quantitative comparisons to be made between conditions where R-loop levels are perturbed. There is increasing interest in factors purported to resolve R-loops in cells (2,6,42), and as many of these factors alter cellular R-loop content, the use of spike-in standards will be particularly important for these studies moving forward.

## Methods

### Cell culture

HeLa cells were obtained from ATCC (Manassas, VA), where they were tested for mycoplasma and verified by STR profiling. These cells were grown in DMEM (Gibco, Dublin, Ireland) supplemented with 10% FBS and 1% penicillin/streptomycin/glutamine, and grown in 5% CO_2_ at 37°C. DRB (Cayman Chemical Company, Ann Arbor, MI) was mixed into pre-warmed media at 100 µM and added to cells for 40 minutes prior to cell harvest.

### Preparation of spike-in hybrids

*E. coli* genomic DNA was prepared as in (https://bio-protocol.org/bio101/e97). Briefly, 1.5 mL of an overnight culture was lysed in 0.6% SDS for one hour, and genomic DNA was extracted using a standard phenol-chloroform isolation. The sequences used for spike-in hybrids were then amplified by PCR using Phusion DNA polymerase, and using primers containing the T7 polymerase promoter sequence. RNA was produced from these PCR templates using *in vitro* transcription from 120 ng of DNA template per spike-in, and purified using an RNeasy kit (#74104, Qiagen, Hilden, Germany), with DNase treatment performed on the columns for 15 min (#79254, Qiagen). The complementary DNA sequence was ordered directly as single-stranded ultramers or megamers (Integrated DNA Technologies, Coralville, IA) (Supplemental Table S1). Annealing was performed between 400 ng of RNA and 100 ng of DNA in 45 µL of 1x buffer 2.1 (#B7202S, New England Biolabs, Ipswich, MA), beginning with a denaturation step for 10 minutes at 95°C, and followed by seventy cycles of 90 second holds at decreasing intervals of 1°C until the temperature reached 25°C. Hybrids were then purified by 1 h of electrophoresis on a 0.9% agarose gel, followed by excision of the hybrid band and purification by centrifugation for 10 minutes at 5000 rpm as in (25). We did not ethanol precipitate the resulting hybrids for cleanup, as we found this caused hybrids to dissociate. We measured the final concentration of nucleic acids after cleanup by fluorimetry using the Qubit HS RNA assay kit (#Q32852, Thermo Fisher Scientific, Waltham, MA). To make the dsDNA spike-ins, sequences were amplified from the *E.coli* genome by PCR using Phusion DNA polymerase, purified by electrophoresis on a 0.9% agarose gel, excised and isolated by gel extraction. The ssDNA spike-in was purchased directly as an ultramer (Integrated DNA Technologies) (Supplemental Table S1). All seven spike-ins were diluted and combined to produce a stock solution of 2.5 pM for each spike-in. The spike-ins were added to individual experimental samples just prior to cell lysis.

### Considerations for spike-in design

In addition to the two synthetic RNA:DNA spike-ins described (∼280 bp long), we also designed, synthesized and purified two additional RNA:DNA spike-ins, each ∼130 bp long. These sequences were successfully synthesized, purified and recovered in DRIP. However, we found that the recommended size-selection with AMPure XP beads (Beckman Coulter, Brea, CA, USA) depleted the smaller spike-ins disproportionately during library preparation. Although altering the size-selection parameters improved the recovery of the shorter spike-ins, this unacceptably compromised library quality. Therefore, care must be taken when using spike-ins <150 bp in length, as these may not be compatible with all library synthesis procedures. Additionally, to provide a significantly cheaper alternative to hybrid spike-in synthesis, we also made hybrids using PCR-generated dsDNA templates for annealing reactions with purified RNA. While we verified that hybrids were successfully made in this way, both DNA strands were present in our sequencing libraries, indicating that some hybrid molecules were formed containing the non-template strand. To generate a more homogeneous hybrid population, we thus proceeded to use hybrids made by annealing RNA to commercially-sourced pure ssDNA templates as described above.

### DRIP

2×10^6^ HeLa cells were lysed by SDS/Proteinase K treatment for 3 h at 37°C. DNA was extracted by phenol-chloroform extraction using phase lock tubes and ethanol precipitated. Precipitated DNA was gently spooled and washed with 70% ethanol without centrifugation. DNA was allowed to air dry for ∼20 minutes and resuspended on ice in 130 uL TE buffer. DNA was sonicated in 6×16 mm microtubes (Covaris, Woburn, MA) to a peak fragment size of 250-300 bp, performed on a Covaris machine (E220 evolution) using SonoLab 7.3 software. For each immunoprecipitation, 4.4 µg of sonicated DNA was used. For RNase H treatment, 8 ug of sonicated DNA was digested with 5 uL RNase H (New England Biolabs) in 1X RNase H digestion buffer overnight at 37°C and purified using standard methods described above. 4 μg of DNA was bound with 10 μg of S9.6 antibody (Antibodies Incorporated, Davis, CA) in 1 X binding buffer (10 mM NaPO4 pH 7, 140 mM NaCl, 0.05% Triton X-100) overnight at 4°C. Dynabeads Protein G beads (Thermo Fisher Scientific) were added for 2 h. The use of magnetic beads strongly increased our fold enrichment compared to agarose beads. Bound beads were washed 3 times in binding buffer and elution was performed in elution buffer (50 mM Tris pH 8, 10 mM EDTA, 0.5% SDS, Proteinase K) for 45 min at 55°C. DNA was purified as described.

### Library preparation and sequencing for qDRIP-seq

DNA libraries were prepared from 3 pooled DRIPs by sample. While the DNA material from one IP (or less) is sufficient for successful library preparation, we found pooling IPs effective in minimizing technical variability due to sample handling in pilot experiments. DNA libraries were synthesized from ssDNA using the Accel-NGS 1S DNA library kit (Swift Biosciences, Ann Arbor, MI), according to the manufacturer’s protocol. Using multiplexing adapters from the 1S Plus Set A Indexing Kit (Swift Biosciences), adapter-ligated DNA was amplified by PCR, then size-selected using a left/right AMPure XP size selection (Beckman Coulter). Library DNA was analyzed on a Bioanalyzer DNA HS (Agilent, Santa Clara, CA) at the Stanford Protein and Nucleic Acid Facility, quantified by qPCR using NEBNext Library Quant Kit for Illumina (New England Biolabs), then pooled and sequenced on a HiSeq 4000 (Illumina, San Diego, CA) at the Stanford Genome Sequencing Service Center, using 2 × 151 bp sequencing.

### qPCR

qPCR was performed on a Roche LightCycler 480 Instrument II using SYBR-Green master mix (Bio-Rad Laboratories, Hercules, CA). Primers used for qPCR are listed in Supplemental Table S2.

### EU nascent transcription assay

Cells were pulsed for 1 hr with 100 μM 5-ethynyl uridine (EU) from the Click-iT RNA Alexa Fluor 488 imaging kit (Thermo Fisher Scientific). Cells were fixed in 4% PFA/PBS for 15 min, and permeabilized with 0.25% Triton/PBS for 15 min. The Click-iT reaction was performed according to manufacturer’s instructions. Cells were then incubated in DAPI for 15 min, and the slides mounted with Pro-Long Gold Antifade (Thermo Fisher Scientific) and imaged on a Zeiss Axioscope with a 20X objective (Zeiss, Oberkochen Germany). EU signal intensity from ≥1200 cells per condition was calculated using CellProfiler (version 3.1.8).

### qDRIP analysis

Prior to alignment, Skewer (43) was used to remove adapter sequences, and Cutadapt (44) was used to remove low complexity G-rich tails from the beginning of R2 with ‘cutadapt -j $N_CORES --minimum-length 30 -U 12’. Trimmed reads were aligned to a custom genome combined from hg38 and the sequences of the synthetic spike-ins with bwa-mem (45). Reads were separated into positive and negative stranded files using SAMtools (46) and unix text-processing utilities. BEDTools (47) was used to convert these SAM files into BEDPE format, with subsequent sorting, filtering for concordant reads, and duplicate alignment removal performed using unix text-processing utilities. Genome browser tracks were produced with the BEDTools genomecov utility, and visualized using IGV (48). Reads from the spike-in did not have duplicate alignments removed, as we expect multiple coincidental fragments to derive from the spike-ins.

### Peak calling and differential peak calling

Peaks were called directly from aligned reads from both strands using MACS2 (49) with default broad peak settings. BEDTools was then used to obtain coverage in each experiment over these consensus peaks. Using these read counts, we filtered out only those regions that showed strong (2-fold) reduction after RNase H treatment as measured by reads per million. We further filtered these peaks to those deriving at least two-thirds of their reads from a single strand.

For differential peak calling, we separately called peaks for the DMSO and DRB samples using the protocol described above. We then used BEDTools to merge these regions in combination with a comparable number of random 5000 bp peaks to include some unchanged background regions to reduce bias in the differential analysis. The DESeq2 R package (50) was then used to obtain fold change estimates and FDR-corrected p-values for differential expression across peaks.

### Metaplots

Metaplots around genes, transcription start sites and other genome features were produced from genome browser tracks with deepTools (51). For gene centered analyses, we used Gencode V29 canonical genes filtered to only include those that had at least one sense read in a publicly available GRO-seq dataset obtained in HeLa cells (35). Tracks for GC and AT-skew were produced using the BEDTools *nuc* tool with further processing from unix text processing tools.

### R-loop absolute quantitation and lifetime analysis

For absolute quantitation, we used BEDTools to calculate maximum read depths over each called peak. We additionally estimated the gene copy number over 10 kb intervals on the genome using our sequenced input samples, finding that most intervals fell into three populations of read-counts with modes having integral ratios of approximately 2:3:4. We thus assigned each 10 kb region as having 2N, 3N or 4N DNA content. Each hybrid peak was assigned a copy number from the modal copy number call across the peak. We then normalized the maximum read depths for peaks in each sample to the estimated genomic copy number at their respective sites, and divided it by the mean spike-in read count obtained for that sample. Per-cell DNA content estimates were done with numerical simulations using NumPy on 200,000 simulated sets of alleles, which were then summed to simulate 100,000 diploid cells. For lifetime analysis, we first used BEDTools to select only those genes with RNase H reversible R-loop peaks. Cell specific rates of transcription were obtained from (37). Using these rates, we estimated the positions of the last uninhibited RNA polymerases after 40 minutes of DRB treatment, and further estimated the time without transcription for 1 kb bins behind these positions. We then calculated spike-in-normalized qDRIP read counts over 500 bp bins over these genes, and took the ratio between the DRB and DMSO signal at each site. Exponential models were fit using linear regression of the natural log of this ratio onto the estimated time since transcription had occurred at each gene. To avoid high leverage points from skewing the estimated regression line, the regression model was weighted by the square root of the time since transcription. Half-lives were calculated by ln(2) divided by the time constant as estimated by regression, for genes well fit by exponential models (Pearson’s R^2^ > 0.8). To calculate the total number of resolution events per day in a cell, we made use of the differential equation for first order decay which relates the rate of decay (dN/dt) to the time constant (k) and the amount of material to decay (N) : dN/dt = -kN. As we found a mean resolution time of 11 minutes, we calculated a mean time constant of ln(2)/(11 minutes) or 0.063/min. Multiplying this by the estimated count of R-loops in a cell at steady state (300), we find that 18.9 R-loops would be resolved per minute, which is approximately 27,000 per day.

### Data processing, data visualization and statistical analysis

Data processing for metaplots, differential peak calls, lifetime analysis, and absolute quantitation were performed with a standard Python scientific stack (Python 3.6.8, NumPy 1.14.0, Pandas 0.22.0). Data was visualized with the Python packages Matplotlib (version 2.1.2) and Seaborn (version 0.8.1). Statistical analysis was primarily performed in Python using the Scipy stats package (1.0.0) for statistical tests and statsmodels (0.8.0) for regression analysis. R (Version 3.1.0) was used for Negative Binomial regression, and for differential peak calling statistics with the Bioconductor DESeq2 package (Love 2014) (version 1.6.3).

## Data Access

All raw and processed sequencing data generated in this study have been submitted to the NCBI Gene Expression Omnibus (GEO; http://www.ncbi.nlm.nih.gov/geo/) under accession number GSE134084.

## Acknowledgments

We thank members of the Cimprich lab for critically reading this manuscript and also Dan Jarosz and Hannah Long for providing helpful feedback. This work was supported by a Fellow Award from the Leukemia and Lymphoma Society to M.P.C. (5455-17), a Stanford Graduate Fellowship and NCI PHS grant (CA09302) to M.J.B., an NIH grant to K.A.C. (GM119334), and a V Foundation Grant (D2018-017) to K.A.C. K.A.C. is an American Cancer Society Research Professor.

## Disclosure Declaration

The authors declare that they do not have any conflicts of interest.

