## Supplementary information for "qDRIP: Quantitative differential RNA:DNA hybrid immunoprecipitation sequencing"

### Supplemental material

Table of contents:

Supplemental figures S1-S7 (2-8)
Supplemental tables S1, S2 (9-13)

Figure S1

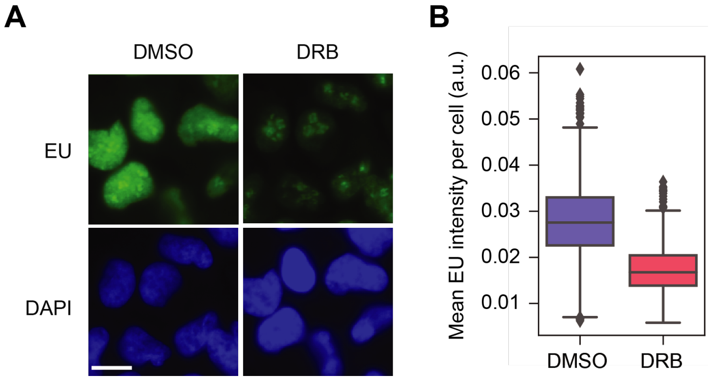

**Figure S1. RNA synthesis measured by EU labelling. (A)** Representative images of nascent EU incorporation in DMSO and DRB treated HeLa cells obtained at 20X magnification. Scale bar is 10 microns. **(B)** Quantification of mean EU incorporation per nucleus in HeLa cells treated with DMSO or DRB ($n \geq 1200$). Box represents the interquartile range of data: whiskers extend 1.5-fold out from the interquartile range, with outliers represented as individual points.

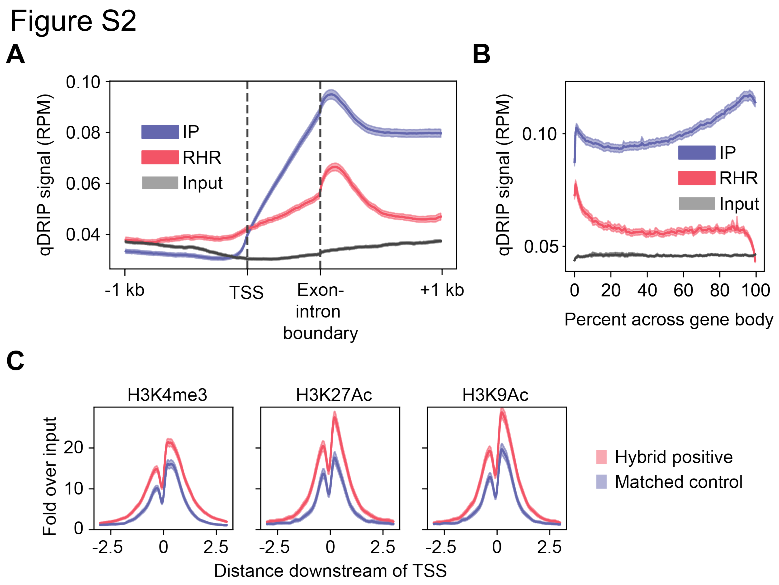

**Figure S2. Additional correlation of genome features to qDRIP signal. (A)** Scaled metaplot of sense hybrids between TSS and first-intron exon boundary, as well as 1 kb upstream of TSS and 1 kb downstream of first intron-exon boundary. Tracks shown are mean IP (blue), RNase H treated IP (red) and pooled input (grey). Bands represent 95% CI of mean read signal. **(B)** Metaplot of S9.6 IP signal over gene bodies. Tracks are same as in (A). **(C)** Distribution of histone marks around TSS of hybrid-containing (red) and expression-matched non-hybrid-containing (blue) genes.

Figure S3

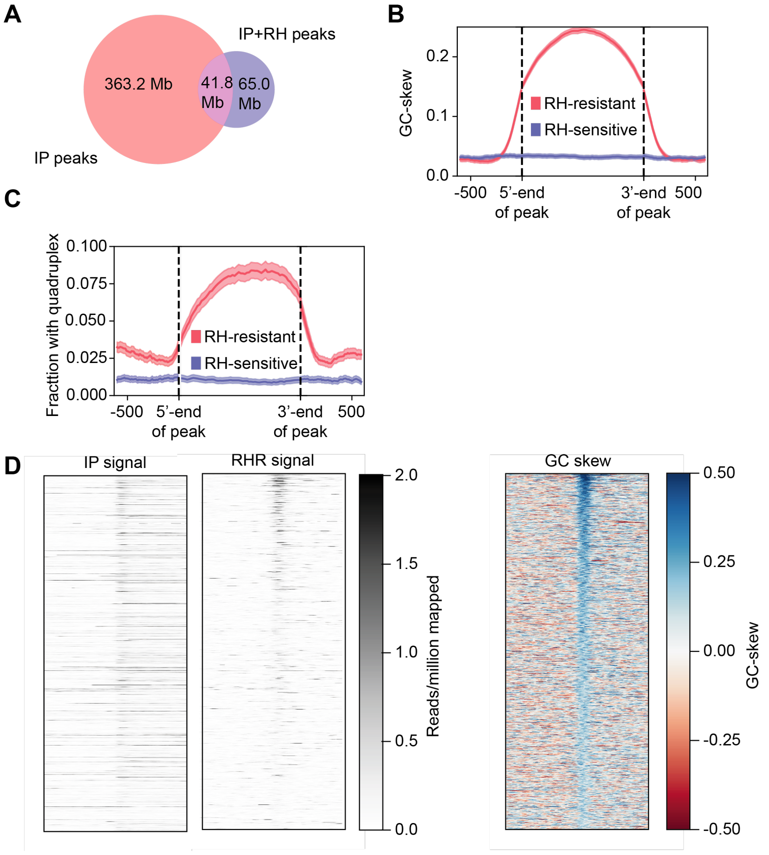

**Figure S3. RNase H insensitive regions have distinct sequence characteristics. (A)** Venn diagram of genome areas (in megabases) occupied by peaks called from qDRIP IP vs input (red), qDRIP RNase H treatment vs input (blue), or their overlap (purple). **(B)** GC-skew around RNase H resistant regions within qDRIP peaks (red) compared to regions of equal lengths randomly selected from non-resistant qDRIP-peaks (blue). Bands represent 95% CI of mean read signal. **(C)** Same as (B), but showing G-quadruplex density over these regions. **(D)** Metaplots of mean IP signal, RNase-H-resistant signal, and GC-skew around top 10000 promoters ranked by GC-skew immediately (0-500 bp) downstream of the TSS.

Figure S4

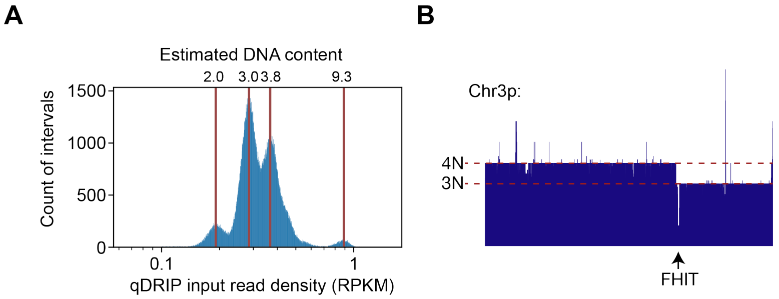

**Figure S4. Copy number estimation in HeLa cells (A)** Histogram of normalized input read counts (as reads per million) across 10 kb bins, showing high variability in genome copy number between different genomic loci. The left cluster of three populations likely represent copy numbers of 2,3, and 4. The rightmost population represents a highly amplified region of chromosome 5p in HeLa cells with approximately 9N DNA content. **(B)** Representative genome browser view showing genome copy number calls over Chr3p. An abrupt copy number change from 4N to 3N content is apparent at the FHIT fragile site.

Figure S5

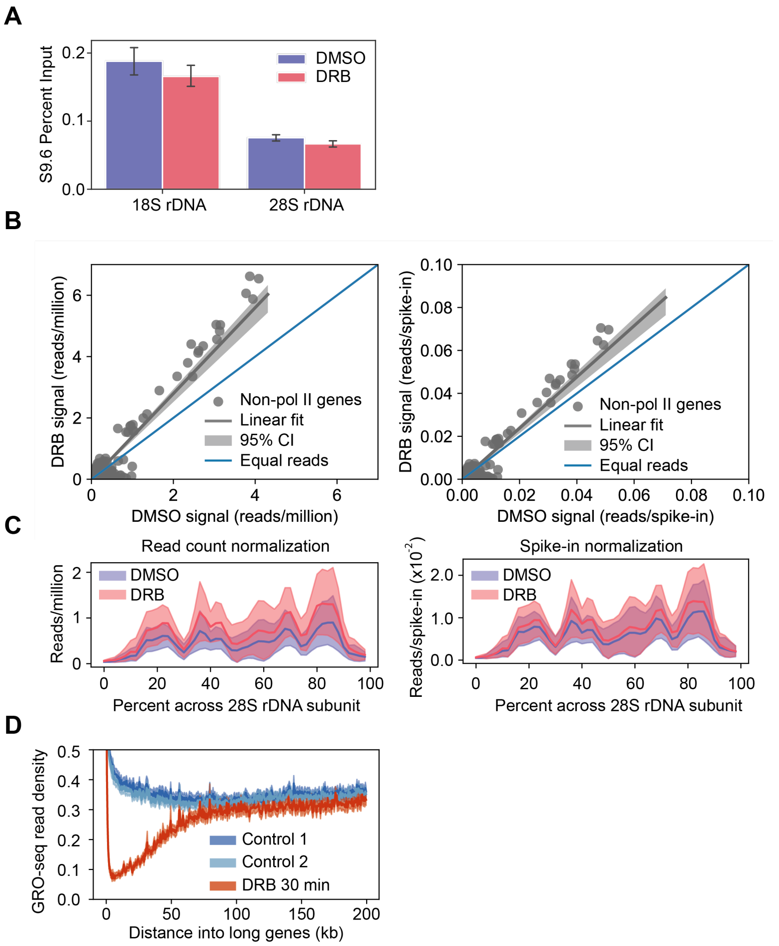

**Figure S5. Validation of differential peak calls. (A)** qPCR measurements of S9.6 IP signal at 18s and 28s rDNA under control (DMSO) and DRB treatment conditions. Error bars represent 95% CI of the mean. **(B)** Scatter plots showing read-counts over Pol I or Pol III-transcribed regions compared between DMSO and DRB using read counts to normalize (left) and spike-ins to normalize (right). Individual regions are shown as grey dots, while the regression line and bootstrapped 95% CI are shown as a grey line and gray band, respectively. Blue diagonal line represents even read counts. The normalization factor calculated using read counts is 1.373 with a bootstrapped 95% CI of (1.247, 1.459), while the normalization factor calculated using spike-ins is 1.170 with a bootstrapped 95% CI of (1.063, 1.242). **(C)** Metaplots of DMSO (blue) and DRB (red) IP signal over Pol I-transcribed 28s rDNA unit normalized using total read counts (left) or mean spike-in counts (right). Bands represent 95% CI of mean read signal. **(D)** Metaplot of nascent transcription as measured by GRO-seq over long genes (>125 kb) from publicly available data (Laitem et al. 2015). Tracks shown are two replicates of DMSO control (dark and light blue) and 100 μM DRB for 30 minutes (red).

Figure S6

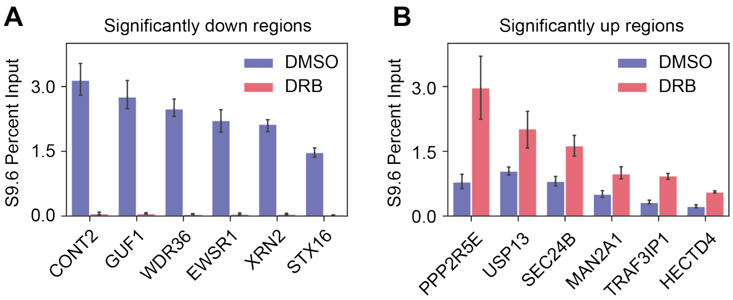

**Figure S6. Validation of spike-in differential peak calls. (A)** qPCR validation of regions called as up in DRB by differential peak calling with spike-ins. (**B)** Same as (A), but for regions that were called as significantly down in DRB by differential peak calling. Error bars represent 95% CI of mean value. All results are significantly different as determined by non-overlapping 95% CIs.

Figure S7

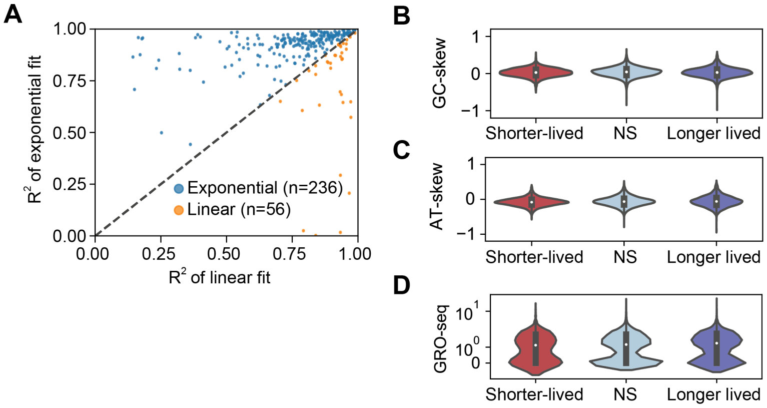

**Figure S7. (A)** Comparison of $R^2$ obtained from exponential and linear fits to DRB/DMSO ratio over estimated time since new transcription. **(B)** G-skew, **(C)** A-skew and **(D)** sense nascent transcription in R-loop regions with shorter than average, close to average, or longer than average lifetimes. (GC-skew p=5.1e-8, AT-skew p=1.8e-6, nascent transcription p=0.001, all by Kruskal-Wallis test).

Table 1: Spike-in sequences

| Name | Sequence | Description |
|---|---|---|
| L132 | CTCATATGCAGCTGGGTGTGTATTTTGTAAACAGAAGTAATTTTCAACTT<br>CTAAGCTTTGTATACAAAGCACTGCCGTAGCAATGCCAAGTGCACGTTT<br>CTTTAATTCAGGAGCGGATTCATACAGGTTGCCCCCTATAGTGAGTCGT<br>ATTACTGCAGGATCC | Template for in vitro transcription |
| H136 | ATAGCCTGCGATGGTGATCAAGTCCGCATAAGCTTTTGCCACGCCAGT<br>CGCGATGGTGCCTACTCCCGGTTCGGAAACCAGCTTCACGGAGATCAT<br>CGCTTTCGGGTTAACCTGCTTGAGGTCGAAAATGAGCTGCCCCTATAGT<br>GAGTCGTATTACTGCAGGATCC | Template for in vitro transcription |
| L286 | ACTGCGACAATAATCCCGGAAATTGACAACCATGCTGGGGCGTTCGGC<br>AGCATAGAAAGCGCCGATAAAATGGTGACCCCTTCTTCTTTAGCAGCTT<br>CAATATACGAAGGTGGAATAGAAAGCAGGCAGCTAAAGACAAAGAACA<br>GTACACTTATGCAGATGATGAGATAAGCGACTTTCATAATTTTTTTGCAT<br>TTATCCATAGCGTGTTCGCCATATTTTTCACGTCTGTCTATGGCAAACGT<br>AGAAATAATGGGCGTATGGCTAAAGGCGAAAACCATCACGGCCCTATA<br>GTGAGTCGTATTACTGCAGGATCC | Template for in vitro transcription |
| H281 | CCAGAACACCATCAACACCCAGCATTCGCGCCGGATGCAGCGACGCCA<br>TATGGATGGCTTCCGCAGGCGTTACGCCCGTCAACTCGACCATATTGC<br>GCACTGCCGCATCAACAGACAGCGTACTGCCCGCCAGCCCACCAGAC<br>GCGGTACGGACAACGCCACCGTGCATCTGCACTTCTTCACCACATAAC<br>GTATAGCGACCATCCGGCATCCCAGCTGCCTGCATCGCGTCGGTGATC<br>AGTACGATTCTCTCTTTCGCACAGCAACAACACAGCGACATTCCCTATA<br>GTGAGTCGTATTACTGCAGGATCC | Template for in vitro transcription |
| L132 | CTCATATGCAGCTGGGTGTGTATTTTGTAAACAGAAGTAATTTTCAACTT<br>CTAAGCTTTGTATACAAAGCACTGCCGTAGCAATGCCAAGTGCACGTTT<br>CTTTAATTCAGGAGCGGATTCATACAGGTTGCC | spike-in sequence and oligo used for RNA:DNA hybrid annealing |
| H136 | ATAGCCTGCGATGGTGATCAAGTCCGCATAAGCTTTTGCCACGCCAGT<br>CGCGATGGTGCCTACTCCCGGTTCGGAAACCAGCTTCACGGAGATCAT<br>CGCTTTCGGGTTAACCTGCTTGAGGTCGAAAATGAGCTGC | spike-in sequence and oligo used for RNA:DNA hybrid annealing |
| L286 | ACTGCGACAATAATCCCGGAAATTGACAACCATGCTGGGGCGTTCGGC<br>AGCATAGAAAGCGCCGATAAAATGGTGACCCCTTCTTCTTTAGCAGCTT<br>CAATATACGAAGGTGGAATAGAAAGCAGGCAGCTAAAGACAAAGAACA<br>GTACACTTATGCAGATGATGAGATAAGCGACTTTCATAATTTTTTTGCAT<br>TTATCCATAGCGTGTTCGCCATATTTTTCACGTCTGTCTATGGCAAACGT<br>AGAAATAATGGGCGTATGGCTAAAGGCGAAAACCATCACGG | spike-in sequence and oligo used for RNA:DNA hybrid annealing |
| H281 | CCAGAACACCATCAACACCCAGCATTCGCGCCGGATGCAGCGACGCCA<br>TATGGATGGCTTCCGCAGGCGTTACGCCCGTCAACTCGACCATATTGC<br>GCACTGCCGCATCAACAGACAGCGTACTGCCCGCCAGCCCACCAGAC<br>GCGGTACGGACAACGCCACCGTGCATCTGCACTTCTTCACCACATAAC<br>GTATAGCGACCATCCGGCATCCCAGCTGCCTGCATCGCGTCGGTGATC<br>AGTACGATTCTCTCTTTCGCACAGCAACAACACAGCGACATT | spike-in sequence and oligo used for RNA:DNA hybrid annealing |

| | | |
|---|---|---|
| LDNA | CATCCCAGAGCGTTGTTGAGTGTAAATGTAGGAGGGAAGATAGTTACA<br>GATCGTGTAATGAGGTCTGTATGGAGTTTCTTTTTTCTTTATACTCTCTT<br>CACGGTGTTTTTTATACTGGTGTTAAATGGTATGGGATATGATTTTCTTA<br>CATCATTTGCAACAGTGGCTGCATGTATTAATAATATGGGATTAGGTTTT<br>GGGGCT | spike-in sequence (genrated by PCR) |
| HDNA | ACTCCGAGCAGCCAGATATCGGAACCGAGCGGCAGCTGTTGCATACGC<br>ATATCCAGCAGGCGCAGATCCAGCGTGCCGTAACGCTGCCACAGCAGC<br>CAGCAAGCAATCGCCAGCAGCAGAGTGCCAAGACGCCCCAGCGCAAA<br>CCACAGTTTGCCCTCTTTGCTGTTGCTGGTGAGGAACACCGCGCACAG<br>GGCCATGATTTCGGCCATTACCACG | spike-in sequence (genrated by PCR) |
| ssDNA | GGCGGTAATGACAGCCAGTACGCCCGGCGCTTTTTGCGCGGCGTCCG<br>TATCAAGGGCGGTGAGGCGTCCTTTGGCAATGGCGGAACCGACGATAT<br>AGCCATAGGCGGCGTTGGGGGCTTCTTCATGCCATTCGTAGGC | spike-in sequence (oligo used directly) |

Table 2: Primer sequences

| Primer name | Sequence | Use |
|---|---|---|
| THUMPD2_F | CAGTGGAGCACTTCTGACTGT | qPCR |
| THUMPD2_R | TGAGTTGGCCATTGGACCTC | qPCR |
| OPN3__F | GGAAGAAAGGGTCTGTCGCA | qPCR |
| OPN3__R | ACCGAACCTGGTTTGCAGAG | qPCR |
| EFNB2__F | TGAGCTTTGCTGTAGGGTGG | qPCR |
| EFNB2__R | AAAGACTTCCACGATGGCGT | qPCR |
| MRPS5__F | TGTTTCAGACCCCGGGAAAG | qPCR |
| MRPS5__R | GCTGTGTTTCACCCGGTTTC | qPCR |
| 18S_F | CGACGACCCATTCGAACGTCT | qPCR |
| 18S_R | CTCTCCGGAATCGAACCCTGA | qPCR |
| 28S_F | AGTCGGGTTGCTTGGGAATGC | qPCR |
| 28S_R | CCCTTACGGTACTTGTTGACT | qPCR |
| SLC30A1__F | CAGCTTTAGTCCTCCTGGGC | qPCR |
| SLC30A1__R | TGTGGGACACTACCACAAGC | qPCR |
| BYSL_F | AAGACTGCCCCTAGTCAGGA | qPCR |
| BYSL_R | ATGCCTGCATGTTTTGGAGC | qPCR |
| CSTF1__F | GACACCGCAGTTCAGTATGC | qPCR |
| CSTF1__R | GGTAGCTACACGGCATGGTC | qPCR |
| SRP19__F | TATACTAATGCTAGGAGAGGAGGG | qPCR |
| SRP19__R | CACAAAAGGACCTACCACAGTTT | qPCR |
| FNDC3A__F | GGATTATGTGAGGTTAGTGGGCT | qPCR |
| FNDC3A__R | ACCCTGTGTAATGAAAACGACAC | qPCR |
| PDE1A__F | CACAAGGTGCTGATGTAGCCA | qPCR |
| PDE1A__R | AAACTCCTTAATTCACCGGGCT | qPCR |
| CACNA2D1__F | CCCCTCACTACCACTACCCA | qPCR |
| CACNA2D1__R | AGGAGAAGAGAGAGCCCAGG | qPCR |
| SBF2_F | ATCAGCAAGCCCAATTCCCA | qPCR |
| SBF2__R | CAAGAAGGGTTTAGTCCCAGGT | qPCR |
| NBAS_F | AGGACAGTGACCGAATCACC | qPCR |
| NBAS_R | ACAAATAGGCCTAAGGCTGGA | qPCR |
| FOXP1__F | ATGACAATCGCAGCCTCACT | qPCR |
| FOXP1__R | GCTTACATAGGGCCGAGCTT | qPCR |
| ATM__F | GAGGAGGTTCTGATCTCACAC | qPCR |
| ATM__R | TCCAAGATCCTGACTTCCTTTGA | qPCR |
| PLXDC2__F | GTGTCAACTGCAAACTCGGG | qPCR |
| PLXDC2__R | TGGGCTAAGCCAGTCTTGTC | qPCR |

| | | |
|---|---|---|
| FSIP1__F | TGCAAGATTGCCCCTGACAT | qPCR |
| FSIP1__R | TGACCCCTTAAGAGGCTGTTG | qPCR |
| KDM6A__F | GGGCCTCCTGTTGTGTTTCT | qPCR |
| KDM6A__R | TATGCGGCAGGGTCAAAGTT | qPCR |
| FAM172A__F | TGGCCCTACTAGAGAACCCC | qPCR |
| FAM172A__R | AGTTCATGAGGCCAGTCTGC | qPCR |
| CONT2__F | GCAGTCATGAGACCTCTATGGT | qPCR |
| CONT2__R | GGCCACAGTGGTAGAAACGA | qPCR |
| GUF1__F | TGTTAGATGTGCCGCTCCAT | qPCR |
| GUF1__R | TTCGTGATGAGGACAACCTGT | qPCR |
| WDR36__F | ACCATCCTAAAGGGCTCTGC | qPCR |
| WDR36__R | TTCTTTAGGAGCTATGCCGCC | qPCR |
| EWRS1__F | TTAGAGAAGATTACAGGCAGACCT | qPCR |
| EWRS1__R | CTTCCGAGCAAGGGAGACTTT | qPCR |
| XRN2__F | CCCTGCTGTCCATCAGGTTT | qPCR |
| XRN2__R | ACCTGCTGGGTTCACAAAGT | qPCR |
| STX16__F | CCCATGCCTCGAGGGTACT | qPCR |
| STX16__R | TTGCCATTACACAATAACCTGCC | qPCR |
| PPP2R5E__F | AATGGCTGATGTGTGGGAACC | qPCR |
| PPP2R5E__R | TTTCCCCTGACACATATCTATTCTG | qPCR |
| USP13__F | CCAGGCTGCTTTCTCCTACC | qPCR |
| USP13__R | ATCCCCGCTGGACTTCCTAT | qPCR |
| SEC24B__F | CTCCTCCTGCCTGCATTCTT | qPCR |
| SEC24B__R | CACACTTCCTGCCCACTTGT | qPCR |
| MAN2A1__F | ATTGCAGTGTCCCTCTGCTG | qPCR |
| MAN2A1__R | AGCATCGTGCCAGAAGCTAA | qPCR |
| TRAF3IP1__F | CCCTGTCTTGGTGGATGGTC | qPCR |
| TRAF3IP1__R | GCTGTATCCGTCCCTCATGG | qPCR |
| HECTD4__F | ATGACTGGTCTTCCCGCTTG | qPCR |
| HECTD4__R | CGCTTCTGAACCTGACACCA | qPCR |
| L132_F | GGATCCTGCAGTAATACGACTCACTATAGGGGGCAACCTGTATGAATCCGC | PCR from E. coli to generate IVT template, and qPCR |
| L132_R | CTCATATGCAGCTGGGTGTG | PCR from E. coli to generate IVT template, and qPCR |
| H136_F | GGATCCTGCAGTAATACGACTCACTATAGGGGCAGCTCATTTTCGACCTCA | PCR from E. coli to generate IVT template, and qPCR |

| | | |
|---|---|---|
| H136_R | ATAGCCTGCGATGGTGATCA | PCR from E. coli to generate IVT template, and qPCR |
| L286_F | GGATCCTGCAGTAATACGACTCACTATAGGGCCGTGATGGTTTTCGCCTTT | PCR from E. coli to generate IVT template, and qPCR |
| L286_R | ACTGCGACAATAATCCCGGA | PCR from E. coli to generate IVT template, and qPCR |
| H281_F | GGATCCTGCAGTAATACGACTCACTATAGGGAATGTCGCTGTGTTGTTGCT | PCR from E. coli to generate IVT template, and qPCR |
| H281_R | CCAGAACACCATCAACACCC | PCR from E. coli to generate IVT template, and qPCR |
| LDNA_F | CATCCCAGAGCGTTGTTGAG | PCR from E. coli to directly generate spike-in and qPCR |
| LDNA_R | AGCCCCAAAACCTAATCCCA | PCR from E. coli to directly generate spike-in and qPCR |
| HDNA_F | ACTCCGAGCAGCCAGATATC | PCR from E. coli to directly generate spike-in and qPCR |
| HDNA_R | CGTGGTAATGGCCGAAATCA | PCR from E. coli to directly generate spike-in and qPCR |
| ssDNA_F | TAATGACAGCCAGTACGCCC | qPCR |
| ssDNA_R | CTACGAATGGCATGAAGAAGC | qPCR |
